# Immunodietica: A data-driven approach to investigate interactions between diet and autoimmune disorders

**DOI:** 10.1101/619700

**Authors:** Iosif M. Gershteyn, Leonardo M.R. Ferreira

**Affiliations:** Ajax Biomedical Foundation, Newton, MA; ImmuVia LLC, Waltham, MA; Department of Surgery, Transplantation Research Laboratory, University of California, San Francisco, San Francisco, CA; Diabetes Center, Sean N. Parker Autoimmunity Research Laboratory, University of California, San Francisco, San Francisco, CA

**Author notes:** These authors contributed equally to this work. To whom correspondence should be addressed. (I.M.G.) or (L.M.R.F.).

**Keywords:** Autoimmunity, diet, epitopes, cross-reactivity, Gershteyn-Ferreira index

## Abstract

Autoimmunity is on the rise around the globe. Diet has been proposed as a risk factor for autoimmunity and shown to modulate the severity of several autoimmune disorders. Yet, the interaction between diet and autoimmunity in humans remains largely unstudied. Here, we systematically interrogated commonly consumed animals and plants for peptide epitopes previously implicated in human autoimmune disease. A total of fourteen species investigated could be divided into three broad categories regarding their content in human autoimmune epitopes, which we represented using a new metric, the Gershteyn-Ferreira index (GF index). Strikingly, pig contains a disproportionately high number of unique autoimmune epitopes compared to all other species analyzed. This work uncovers a potential new link between pork consumption and autoimmunity in humans and lays the foundation for future studies on the impact of diet on the pathogenesis and progression of autoimmune disorders.

## 1. Introduction

Autoimmune disorders, combined, affect an estimated 50 million Americans. The National Institutes of Health (NIH) spend over $500M annually in research grants to study autoimmunity. Yet, the risk factors, causes and modifiers of most autoimmune disorders remain poorly understood. Strikingly, genetics has been estimated to account for only 30% of all autoimmune diseases, with environmental triggers being responsible for the vast majority of autoimmune disease incidence (Bach, 2002; Vojdani et al., 2014).

### 1.1. Molecular mimicry

The most well known example of an autoimmunity environmental trigger is infection. Some autoimmune disorders, notably Guillain-Barré syndrome and type 1 diabetes (T1D), have been suggested to be triggered by bacterial and viral infections (Menser et al., 1978; Schopfer et al., 1982; Lonnrot et al., 2000; Shahrizaila and Yuki, 2011). The mechanism behind these observations is thought to be molecular mimicry. Some bacteria and viruses contain epitopes identical to self epitopes in the host. While most self-reactive T cells are deleted in thymus (central tolerance), some escape to the periphery and can be detected in peripheral blood (Yu et al., 2015). Nevertheless, these autoreactive T cells remain anergic in the absence of danger signals and thus pose no threat of autoimmunity at steady. Infection with bacteria or viruses containing these cross-reactive peptides, however, will lead the host to mount an immune response against them potentially using T cells that recognize pathogenic antigens that match self antigens. When infection subsides, the remaining memory T cells that recognize those shared bacterial and viral antigens are now poised to attack the tissues that express their cognate antigens, leading to autoimmunity (Guilherme et al., 1995; Rojas et al., 2018).

A decade ago, a dramatic demonstration that exposure to higher organisms’ antigens can also cause autoimmunity occurred when workers processing pig brains in a pork plant suddenly developed an autoimmune neurologic disorder. It was later determined that exposure to aerosolized pig neural antigens induced polyradiculoneuropathy in as little as 4 weeks of exposure, with neural-reactive IgG antibodies being detected in the serum of all exposed workers (Lachance et al., 2010; Tracy and Dyck, 2011).

### 1.2. Oral tolerance and gut permeability

Can dietary antigens then initiate or exacerbate autoimmune disease? Oral antigen delivery often results in the creation of tolerance to that antigen, a phenomenon known as oral tolerance. Initially described in the mouse, oral tolerance is induced upon the interaction of orally administered antigens with the immune system in the gut, chiefly in the mesenteric lymph nodes, leading to the generation of antigen-specific regulatory T cells (Chen et al., 1994; Pabst and Mowat, 2012; Esterhazy et al., 2016). Yet, oral tolerance can be compromised. Recently, it was found that mechanical injury to the skin promotes anaphylaxis, an acute allergic reaction, to oral antigen. Mechanistically, injury-activated keratinocytes in the skin released IL-33 systemically, ultimately activating gut mast cells and increasing gut permeability (Leyva-Castillo et al., 2019). Intermittent gut permeability allows partially digested foreign proteins and commensal bacteria to enter the bloodstream and be presented to the immune system in a pro-inflammatory context. Recognition of cross-reactive diet-derived peptides by autoreactive T cells in this fashion may initiate or help sustain autoimmunity (Bischoff et al., 2014; Mu et al., 2017).

### 1.3. Diet and autoimmune disease

Indeed, several instances of an impact of diet on autoimmunity have been reported. Rheumatoid arthritis (RA) patients often report an association between food intake and RA symptom severity. In line with this observation, all three immunoglobulin classes (IgG, IgA, and IgM) were found to have heightened food-specific activities both systemically (serum) and in the intestine (jejunal perfusion fluid) in most of the RA patients analyzed as compared to healthy controls. Recognized antigens originated from cow’s milk, cereals, hen’s eggs, cod, and pork. The authors hypothesized the occurrence of immune complexes in the joints, as a result of many modest hypersensitivity reactions to food antigens, as a mechanism to explain the patients’ reported discomfort (Hvatum et al., 2006). Remarkably, a strong correlation (r^2^ = 0.795) was found between cow milk consumption and multiple sclerosis (MS) incidence across 27 countries. Of note, no such correlation was found with cheese consumption, suggesting the involvement of molecules not present in processed dairy (Malosse and Perron, 1993). Of note, introducing certain foods early in life has been associated with T1D development. Yet, studies in this field have yielded contradictory results, warranting the use of bigger cohorts and more mechanistic studies in the future (Kostraba et al., 1993; Norris et al., 2003; Virtanen and Knip, 2003).

Porcine neural antigens contain epitopes recognized in human neural autoimmunity (Lachance et al., 2010; Tracy and Dyck, 2011). Are there porcine peptide epitopes recognized in autoimmune diseases in other tissues? More broadly, what is the degree of similarity between common food epitopes and those implicated in human autoimmune disorders? Are specific animals or plants enriched for cross-reactive epitopes in certain diseases? We sought to systematically address these questions by mapping the epitope similarity between animals and plants commonly consumed and different human autoimmune diseases.

## 2. Materials and methods

To investigate the overlap of epitopes between human autoimmune disease and commonly consumed foods, we aggregated all epitopes implicated in 77 diseases available at www.iedb.org and cross-referenced them with every epitope in 14 organisms. We utilized Node.js, AWS EC2, and PostgreSQL database technologies in order to automate data gathering and allow for expedient analysis. Data gathering automation was broken into two custom built processes running concurrently

### 2.1. Process #1

The first process (Process #1) queried the National Center for Biotechnology Information (NCBI) blastp system. The query request consisted of the unique epitope ID and the list of organisms that were considered for the study. All the queries went against the NCBI refseq_protein database. Initially, the process made a query for a single epitope and a single organism. However, this approach would not scale for our use case, as the final data set would consist of ∼18000 epitopes in combination with 14 organisms (in excess of 250,000 queries). Additionally, depending on the query and its position in the job queue on the NCBI servers, each result would vary in time spanning from under a minute to multiple hours. Fortunately, the API supports a list of organisms as a valid query criterion, significantly cutting down the round trip of each query. Once a request was made for a particular epitope in the database, Process #1 created a record with the Request ID where the uniqueness of the record was the epitope ID and the organism. The record was marked as pending to indicate that it is ready to be retrieved for processing by Process #2. The process continued until all the epitopes had been queried and marked as pending.

### 2.2. Process #2

The second process (Process #2) retrieved the results of the queries made by using the Request ID from Process #1. As the query could take a certain amount of time to finish, the process would iterate through the list of all the records that had a valid Request ID to obtain the results. In the query result data, the top hit for each organism was selected and then, if the Query Coverage and Identity Percentage values were both 100, the record result was considered as a hit (total match), otherwise a miss. The record was marked as complete so the process would ignore it on the next pass. The system continued polling NCBI servers until all results were gathered.

Once both processes were fully executed, we had compiled a fully comprehensive database of autoimmune human-organism epitope overlap. An interactive website for researchers and the public to explore this database will be available at www.immunodietica.com.

## 3. Results

After screening all available linear epitopes of a total of 14 commonly consumed animals and plants against human linear epitopes implicated in autoimmune disorders, we selected hits with 100% coverage and identity (identical amino acid sequences) and rank ordered the hits for immunogenic similarity (see *Materials and Methods*). We then normalized this index, which we named the Gershteyn-Ferreira index (GF index), to range from 0 to 1. As can be seen in Figure 1, the analyzed species fell within 3 major groups with regards to their global GF index. Red meat (cow, sheep, goat, pig) had the highest GF index, poultry and fish (chicken, turkey, duck, tilapia, and salmon) an intermediate GF index, and cereals (rice, quinoa, soybean, rye, and wheat) a low GF index.

**Figure 1.**
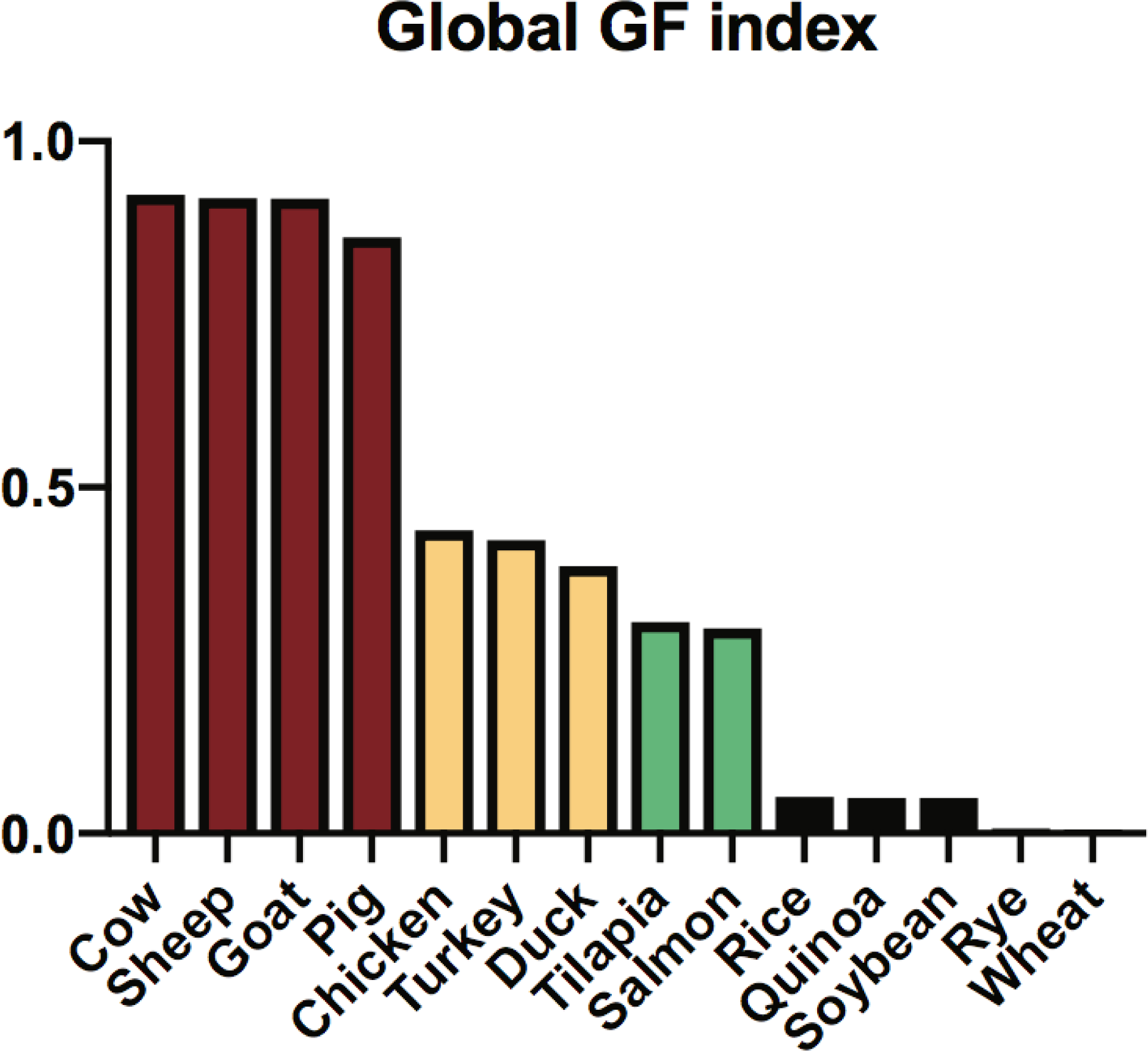
Global Gershteyn-Ferreira (GF) index for different species. Species are colored according to food group: red meat, poultry, fish, and cereals. The global GF index ranges between 0 and 1. Rye and wheat have very low non-zero GF indexes.

In order to dissect the relative similarity of each dietary epitope to specific autoimmune disorders, we isolated autoimmune disease-associated epitopes and cross-referenced each organism’s hits with the overlap to each of the 77 immune diseases featured in the www.iedb.org database (Figure 2).

**Figure 2.**
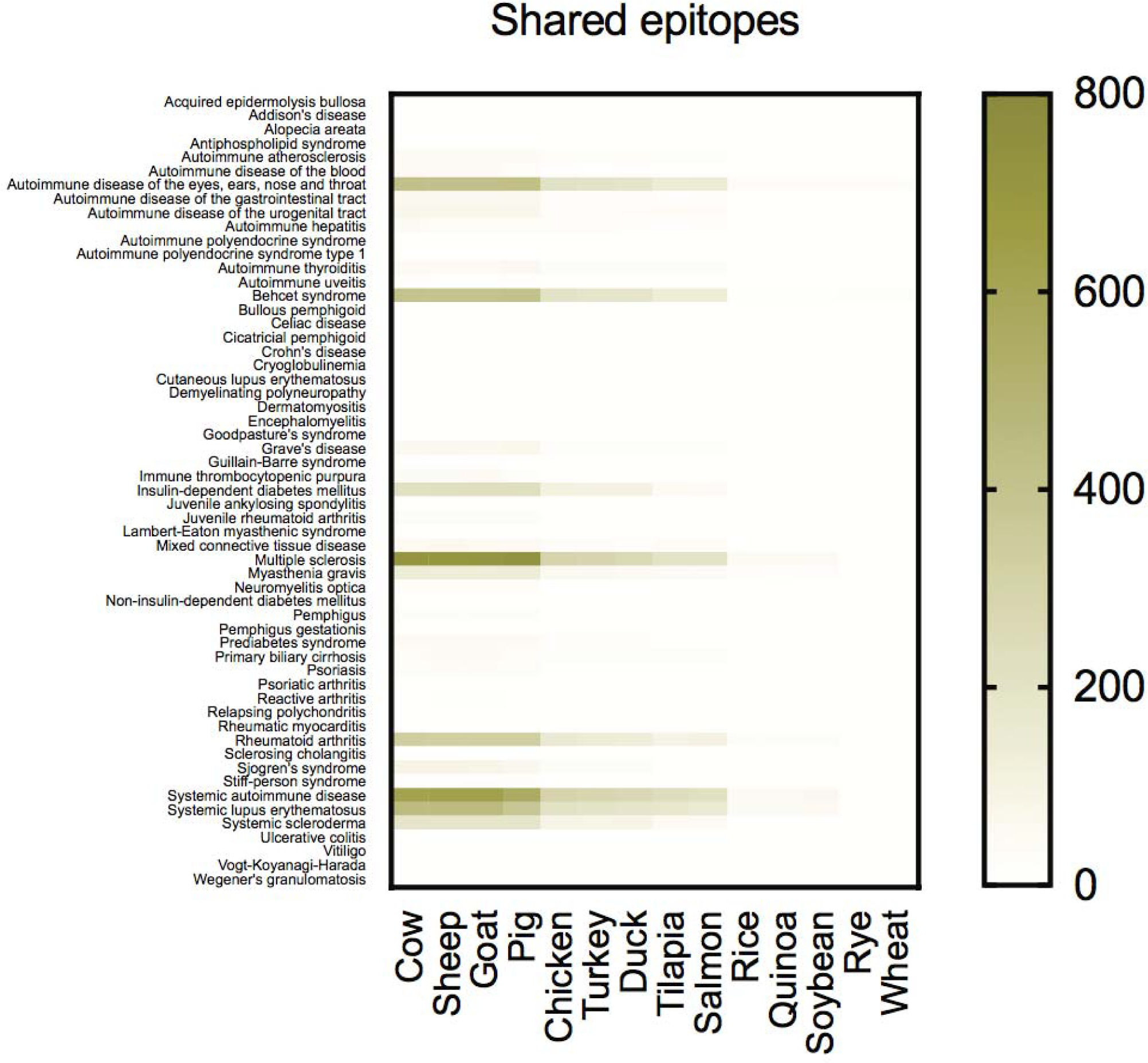
Shared epitopes per autoimmune disease. Autoimmune diseases are listed on the y axis and analyzed species on the x axis.

Similarly to the global GF index, cow, sheep, goat, pig (red meat) had the highest number of epitopes shared with specific autoimmune diseases (Figure 2). The highest number of shared epitopes was 725; pig has 725 epitopes implicated in MS.

Finally, we isolated the unique hits per animal or plant and found that the amount of unique disease-specific epitopes appearing in pigs is 10-15 times higher than in any other organism (Figure 3). Of the 68 diseases analyzed (9 diseases were excluded due to the lack of dietary antigen hits), 40 have at least one unique autoimmune epitope found in one of the animals and plants analyzed (Figure 3).

**Figure 3.**
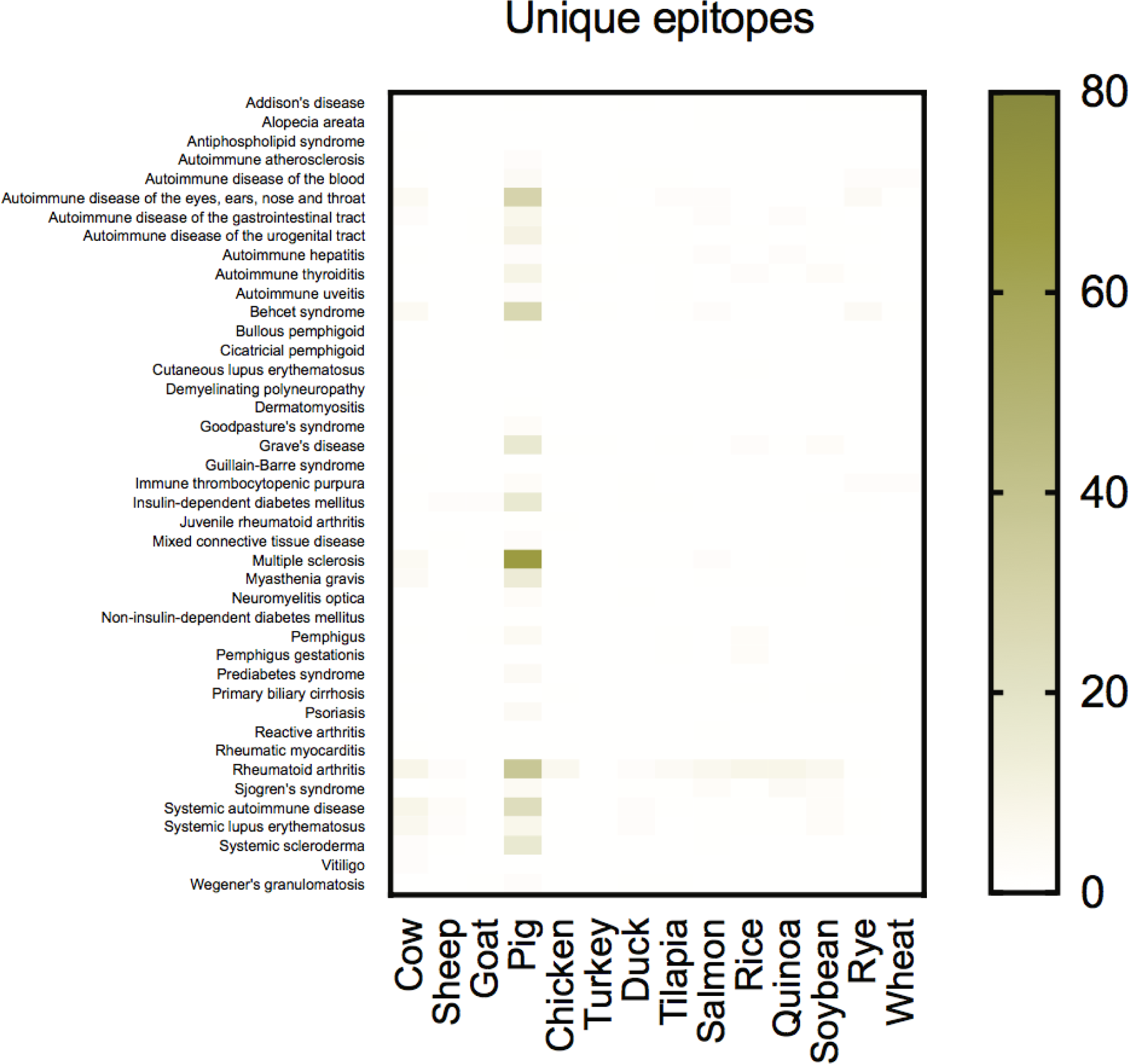
Unique epitopes per autoimmune disease. Autoimmune diseases are listed on the y axis and analyzed species on the x axis.

Strikingly, pig was found to share at least one epitope with 29 of those diseases (72.5%). Thirteen diseases only had epitope matches with one species (Table 1). Pig shared unique epitopes with 4 (30.8%) of these disorders: autoimmune atherosclerosis, bullous pemphigoid, dermatomyositis, and Goodpasture’s syndrome. Yet, the species found to possess shared unique epitopes with the highest number of diseases (5) in exclusivity was the cow. These were APS, demyelinating polyneuropathy, Guillain-Barre syndrome, rheumatic myocarditis, and vitiligo (Table 1).

**Table 1.** Diseases with single species epitope matches

| **Disease** | **Species** | **Number of hits** |
| --- | --- | --- |
| Alopecia aerata | Salmon | 1 |
| Antiphospholipid syndrome | Cow | 1 |
| Autoimmune atherosclerosis | Pig | 2 |
| Bullous pemphigoid | Pig | 1 |
| Cutaneous lupus erythematosus | Rice | 1 |
| Demyelinating polyneuropathy | Cow | 1 |
| Dermatomyositis | Pig | 1 |
| Goodpasture's syndrome | Pig | 3 |
| Guillain-Barre syndrome | Cow | 1 |
| Non-insulin dependent diabetes mellitus | Rye | 1 |
| Reactive arthritis | Salmon | 1 |
| Rheumatic myocarditis | Cow | 1 |
| Vitiligo | Cow | 2 |

To better illustrate the unique links between peptide epitopes present in various commonly consumed plants and animals and autoimmune disorders, we created the Autoimmune unique GF index, where we normalized the unique epitope matches to the maximum number observed, i.e. 325 unique epitope matches found in pig (Figure 4).

**Figure 4.**
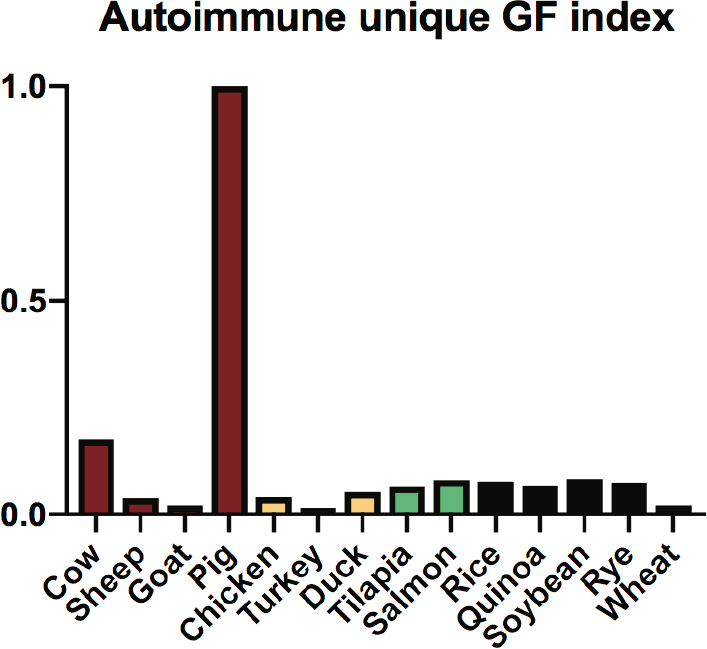
Autoimmune unique Gershteyn-Ferreira (GF) index. Species are colored according to food group: red meat, poultry, fish, and cereals. The autoimmune unique GF index ranges between 0 and 1.

## 4. Discussion

Diet has undergone tremendous changes over the course of history due to globalization and increased life standards. In particular, the consumption of meat and animal products in general has increased over the past decades in both the developed and the developing world (Speedy, 2003). The same time period has also witnessed a noted rise in autoimmune disorders, overwhelmingly due to environmental factors (Bach, 2002). In addition to many other environmental factors tested, consumption of specific foods has been associated with the genesis and severity of autoimmune disease (Malosse and Perron, 1993; Hvatum et al., 2006). Through systematic analysis, we found many peptide epitopes present in species commonly consumed as food that have been implicated in human autoimmune disorders. Of note, pig contained the highest number of autoimmune-associated epitopes not found in any other species investigated.

A caveat in our current approach is the inability to account for immunogenic peptides arising from protein posttranslational modifications inside cells. Three examples stand out: gliadin peptides modified by tissue transglutaminase involved in the pathogenesis of celiac disease (Molberg et al., 1998), ‘hybrid insulin peptides” (HIPs), fusion peptides resulting from covalent cross-linking between pro-insulin peptides and peptides derived from other proteins present in β cells’ secretory granules recently implicated in T1D (Delong et al., 2016), and citrullinated proteins, thought to be important targets for autoantibody recognition in RA (Mathsson et al., 2008).

We predict that subpopulations with outsized contributions of certain foods in their diet where there is a shared epitope with an autoimmune disease will be over-represented in the incidence of that disease in the total population, assuming their genetic predisposition is not significantly different from the average in that population. Moreover, avoidance of foods containing autoimmune peptide epitopes could improve ongoing symptoms of the autoimmune disease in question.

Interestingly, many cultures forbid certain foods from consumption. Pork specifically has been considered unclean by a vast variety of cultures and religions. Since these rules have stood the test of time and are likely to remain for many years going forward, it is intriguing that we found pig antigens to contain, by far, the highest number of unique overlapping epitopes with human autoimmune disorder-associated epitopes. Our results support the hypothesis that there are links between specific foods and autoimmune diseases, and that a common denominator is disproportionately porcine in nature.

## Acknowledgements

The authors thank Andrey Burov for his excellent technical and programming contributions, and Dr. Andrey Vyshedskiy for his stimulating discussions. This work was supported by the Ajax Biomedical Foundation (Newton, MA). L.M.R.F. is a Jeffrey G. Klein Family Diabetes Fellow. Dedicated to the memory of Lev Gershteyn.

## References

Bach, J.F., 2002. The effect of infections on susceptibility to autoimmune and allergic diseases. N Engl J Med 347, 911–20.

Bischoff, S.C., Barbara, G., Buurman, W., Ockhuizen, T., Schulzke, J.D., Serino, M., Tilg, H., Watson, A. and Wells, J.M., 2014. Intestinal permeability--a new target for disease prevention and therapy. BMC Gastroenterol 14, 189.

Chen, Y., Kuchroo, V.K., Inobe, J., Hafler, D.A. and Weiner, H.L., 1994. Regulatory T cell clones induced by oral tolerance: suppression of autoimmune encephalomyelitis. Science 265, 1237–40.

Delong, T., Wiles, T.A., Baker, R.L., Bradley, B., Barbour, G., Reisdorph, R., Armstrong, M., Powell, R.L., Reisdorph, N., Kumar, N., Elso, C.M., DeNicola, M., Bottino, R., Powers, A.C., Harlan, D.M., Kent, S.C., Mannering, S.I. and Haskins, K., 2016. Pathogenic CD4 T cells in type 1 diabetes recognize epitopes formed by peptide fusion. Science 351, 711–4.

Esterhazy, D., Loschko, J., London, M., Jove, V., Oliveira, T.Y. and Mucida, D., 2016. Classical dendritic cells are required for dietary antigen-mediated induction of peripheral T(reg) cells and tolerance. Nat Immunol 17, 545–55.

Guilherme, L., Cunha-Neto, E., Coelho, V., Snitcowsky, R., Pomerantzeff, P.M., Assis, R.V., Pedra, F., Neumann, J., Goldberg, A., Patarroyo, M.E. and et al., 1995. Human heart-infiltrating T-cell clones from rheumatic heart disease patients recognize both streptococcal and cardiac proteins. Circulation 92, 415–20.

Hvatum, M., Kanerud, L., Hallgren, R. and Brandtzaeg, P., 2006. The gut-joint axis: cross reactive food antibodies in rheumatoid arthritis. Gut 55, 1240–7.

Kostraba, J.N., Cruickshanks, K.J., Lawler-Heavner, J., Jobim, L.F., Rewers, M.J., Gay, E.C., Chase, H.P., Klingensmith, G. and Hamman, R.F., 1993. Early exposure to cow’s milk and solid foods in infancy, genetic predisposition, and risk of IDDM. Diabetes 42, 288–95.

Lachance, D.H., Lennon, V.A., Pittock, S.J., Tracy, J.A., Krecke, K.N., Amrami, K.K., Poeschla, E.M., Orenstein, R., Scheithauer, B.W., Sejvar, J.J., Holzbauer, S., Devries, A.S. and Dyck, P.J., 2010. An outbreak of neurological autoimmunity with polyradiculoneuropathy in workers exposed to aerosolised porcine neural tissue: a descriptive study. Lancet Neurol 9, 55–66.

Leyva-Castillo, J., Galand, C., Kam, C., Burton, O., Ziegler, S.F., Chiu, I., Austen, F. and Geha, R.S., 2019. Mechanical Skin Injury Promotes Food Anaphylaxis by Driving Intestinal Mast Cell Expansion. Immunity 50, 1–14.

Lonnrot, M., Korpela, K., Knip, M., Ilonen, J., Simell, O., Korhonen, S., Savola, K., Muona, P., Simell, T., Koskela, P. and Hyoty, H., 2000. Enterovirus infection as a risk factor for beta-cell autoimmunity in a prospectively observed birth cohort: the Finnish Diabetes Prediction and Prevention Study. Diabetes 49, 1314–8.

Malosse, D. and Perron, H., 1993. Correlation analysis between bovine populations, other farm animals, house pets, and multiple sclerosis prevalence. Neuroepidemiology 12, 15–27.

Mathsson, L., Mullazehi, M., Wick, M.C., Sjoberg, O., van Vollenhoven, R., Klareskog, L. and Ronnelid, J., 2008. Antibodies against citrullinated vimentin in rheumatoid arthritis: higher sensitivity and extended prognostic value concerning future radiographic progression as compared with antibodies against cyclic citrullinated peptides. Arthritis Rheum 58, 36–45.

Menser, M.A., Forrest, J.M. and Bransby, R.D., 1978. Rubella infection and diabetes mellitus. Lancet 1, 57–60.

Molberg, O., McAdam, S.N., Korner, R., Quarsten, H., Kristiansen, C., Madsen, L., Fugger, L., Scott, H., Noren, O., Roepstorff, P., Lundin, K.E., Sjostrom, H. and Sollid, L.M., 1998. Tissue transglutaminase selectively modifies gliadin peptides that are recognized by gut-derived T cells in celiac disease. Nat Med 4, 713–7.

Mu, Q., Kirby, J., Reilly, C.M. and Luo, X.M., 2017. Leaky Gut As a Danger Signal for Autoimmune Diseases. Front Immunol 8, 598.

Norris, J.M., Barriga, K., Klingensmith, G., Hoffman, M., Eisenbarth, G.S., Erlich, H.A. and Rewers, M., 2003. Timing of initial cereal exposure in infancy and risk of islet autoimmunity. JAMA 290, 1713–20.

Pabst, O. and Mowat, A.M., 2012. Oral tolerance to food protein. Mucosal Immunol 5, 232–9.

Rojas, M., Restrepo-Jimenez, P., Monsalve, D.M., Pacheco, Y., Acosta-Ampudia, Y., Ramirez-Santana, C., Leung, P.S.C., Ansari, A.A., Gershwin, M.E. and Anaya, J.M., 2018. Molecular mimicry and autoimmunity. J Autoimmun 95, 100–123.

Schopfer, K., Matter, L., Flueler, U. and Werder, E., 1982. Diabetes mellitus, endocrine autoantibodies, and prenatal rubella infection. Lancet 2, 159.

Shahrizaila, N. and Yuki, N., 2011. Guillain-barre syndrome animal model: the first proof of molecular mimicry in human autoimmune disorder. J Biomed Biotechnol 2011, 829129.

Speedy, A.W., 2003. Global production and consumption of animal source foods. J Nutr 133, 4048S–4053S.

Tracy, J.A. and Dyck, P.J., 2011. Auto-immune polyradiculoneuropathy and a novel IgG biomarker in workers exposed to aerosolized porcine brain. J Peripher Nerv Syst 16 Suppl 1, 34–7.

Virtanen, S.M. and Knip, M., 2003. Nutritional risk predictors of beta cell autoimmunity and type 1 diabetes at a young age. Am J Clin Nutr 78, 1053–67.

Vojdani, A., Pollard, K.M. and Campbell, A.W., 2014. Environmental triggers and autoimmunity. Autoimmune Dis 2014, 798029.

Yu, W., Jiang, N., Ebert, P.J., Kidd, B.A., Muller, S., Lund, P.J., Juang, J., Adachi, K., Tse, T., Birnbaum, M.E., Newell, E.W., Wilson, D.M., Grotenbreg, G.M., Valitutti, S., Quake, S.R. and Davis, M.M., 2015. Clonal Deletion Prunes but Does Not Eliminate Self-Specific alphabeta CD8(+) T Lymphocytes. Immunity 42, 929–41.

